# The Anti-inflammatory Drug Leflunomide Inhibits NS2B3 Cluster Formation During Dengue Viral Infection as Revealed by Single Molecule Imaging

**DOI:** 10.1101/2023.12.26.573168

**Authors:** Jiby Mary Varghese, S Aravinth, Neeraj Pant, Partha Pratim Mondal

## Abstract

A prerequisite for Dengue viral infection is the clustering of NS2B3 viral protein in the infected cell. This calls for drugs capable of reversing the biological processes leading to the declustering of NS2B3 viral complex. In this work, we report a new drug (leflunomide) that shows reversal of NS2B3 clustering, post 24 hours of cell transfection with a recombinant probe (Dendra2-NS2B3) containing the viral complex of interest (NS2B3). To study, we constructed a photoactivable recombinant plasmid for visualizing the activity of the target protein-of-interest (Dendra2-NS2B3). This enabled a better understanding of the underlying biological processes involved in Dengue and the role of NS2B3. The study was performed in a cellular system by transfecting the cell (NIH3T3 -mouse fibroblast cell line), followed by drug treatment studies. A range of physiologically relevant concentrations (250 *nM −* 10 *μM*) of the FDA-approved drug (leflunomide) was used. The single molecule super-resolution microscopy (*scanSMLM*) study showed declustering of NS2B3 clusters for concentrations *>* 250 *nM* and near complete disappearance of clusters at concentrations *>* 5 *μM* . Moreover, the associated critical biophysical parameters suggest a substantial decrease in clustered molecules (from 53.2 *±* 1.77% for control to 14.89 *±* 4.80% at 250 *nM*, and further reduction to 10.55 *±* 2.91% at 500 *nM*). Moreover, the number of clusters reduced from 46 *±* 15 to 13 *±* 4, and the number of molecules per cluster decreased from 133 *±* 29 to 62 *±* 3, with a depletion in large clusters (from 24 to 12). The parameters collectively indicate the clustering nature of NS2B3 viral protein during the infection process at a cellular level and the effect of leflunomide in declustering. The results supported by statistical analysis suggest strong declustering promoted by leflunomide, which holds the promise to contain/treat dengue viral infection.

**Statement of Significance:** The fact that there is no approved antiviral approach for Dengue makes it life-threatening and calls for ways to tackle viral infection. Hence, understanding Dengue biology at a single molecule level plays a vital role. In the present super-resolution study, we noted the formation of key viral protein (NS2B3) clusters post 24 hours of transfection in a cellular system. We identified a repurposed FDA-approved drug (Leflunomide) that inhibits the clustering process and promotes declustering at higher drug concentrations. This may become the basis of future studies, which may have therapeutic potential against Dengue.

## I. INTRODUCTION

Dengue fever is a mosquito-borne disease caused by Dengue virus of family Flaviviridae. It is endemic in 110 countries and approximately 400 million people get infected every year with a death rate of 20,000 annually [1]. Dengue viral genome carries a positive single-stranded RNA, which encodes a polyprotein that ultimately produces 3 structural proteins (capsid, membrane precursor (prM), envelope) and 7 nonstructural proteins NS1, NS2A, NS2B, NS3, NS4A, NS4B, and NS5 [2]. Out of nonstructural proteins, NS3 is a multifunctional protein with protease activity along with NS2B as a co-factor. Together the complex is known as NS2B3 protease. The NS2B3 protease complex holds special attention as it possesses proteolytic activity and processing of viral polyprotein at the junctions with other nonstructural proteins (capsid-prM, NS2A/NS2B, NS2B/NS3, NS3/NS4A, NS4B/NS5, internal NS2A, NS3, and NS4A)[3] [4]. These proteolytic cleavages are essential for viral particle maturation and release [5]. Apart from its proteolytic activity on viral polyprotein, NS2B3 protease interacts with host subcellular organelles to create a favorable viral invasion environment. Reports suggest that NS3 protein interacts with fatty acid synthase of endoplasmic reticulum leading to increased fatty acid synthesis and thereby efficient viral packaging [6]. The NS2B3 protease also targets mitochondria by inducing cleavage to matrix-localized GrpE protein homolog 1 (GrpEL1) of mitochondria [7]. It also cleaves mitochondrial outer membrane protein mitofusin 1 and 2 (MFN 1/2), leading to mitochondrial fragmentation and dengue-induced cell death[8]. Apart from these facts, NS2B3 protein sequence is sufficiently conserved across all four serotypes of dengue virus (Zika, West-Nile, Encephalitis, Yellow-fever). The multifunctionality and conserved genetic sequence make NS2B3 protease a potential drug target.

As viruses are very small structures usually *<* 200*nm*, conventional microscopic techniques are not capable of resolving them and their related polyproteins. To overcome this, single-molecule super-resolution microscopes are ideal since it can resolve features beyond the diffraction limit (*∼ λ/*2). Of all the existing super-resolution techniques (such as STED, structured illumination, and its recent variants), fluorescence photoactivation localization microscopy (FPALM) and scanning SMLM (*scanSMLM*) holds promise since it can quantify individual molecules and their collective dynamics with high temporal resolution in a cell volume [24][12]. The technique uses photoactivable fluorescent probes that is activated by a high-frequency laser and convert a small fraction of inactive fluorophore to active form. Subsequently, the activated molecules are excited by another light and the fluorescence is collected by a sensitive detector (EMCCD / sCMOS). Using dedicated computational process, the bright spots / blobs in the recorded frame can be localized. Post emission, the molecules may get photobleached. Similarly, remaining inactive fluorophores also repeat the same process until sufficient molecules have been localized. This produces a super-resolved image of the specimen, revealing the location of fluorescent probes. When conjugated with the protein-of-interest, the technique reveals the location of single molecules and their interactions with the inter-cellular organelles [24].

In general, viral proteins are known to form clusters in the localized region of the cell. In Influenza, the clustering of hemagglutinin protein is well-known and studied extensively [9] [10]. Drug repurposing is a strategy for identifying new therapeutic uses for an already existing FDA-approved drug. In the present study, we investigate a new role of an already FDA-approved drug that can interfere with the clustering of NS2B3 and may prove to have therapeutic potential against dengue. We have chosen an antiinflammatory drug leflunomide, which is known to induce mitochondrial fusion by activating mitofusin 1 and 2 [11]. For the first time, we study the interaction of NS2B3 protease-induced changes in the cellular system at the single-molecule level and investigate whether leflunomide can reverse the clustering effect induced by NS2B3 viral protein [17]. This work explores its potential to induce declustering through direct or indirect mechanisms. We use single-molecule super-resolution *scanSMLM* microscopy to reveal the action of leflunomide against NS2B3-induced changes (clustering process) at single molecule level. Although leflunomide has shown other activities (such as mitochondrial fusion), we observed its declustering behavior for the present study related to NS2B3 protein clustering during Dengue infection. The study is carried out at a cellular level by transfection process. The mechanism of action may be direct on NS2B3 cluster or indirect via some other processes involving organelles such as mitochondria. The exact mode of action needs to be investigated further, but we realize the therapeutic potential of leflunomide in reversing the action induced by key NS2B3 viral protein.

Through this study, we hope to select drug targets to disrupt the onset of the clustering process, hoping to inhibit the chain of viral infection.

## II. RESULTS

### A. Construction of photoactivable recombinant probe and its validation using confocal microscopy

The first step in the proposed study requires the construction of a photoactivable probe (Dendra2) that can be conjugated with the protein of interest (here, NS2B3). The resultant recombinant probe can be used to visualize the collective dynamics of protein-of-interest and its interactions with sub-cellular organelles. Sub-cloning strategies were developed and executed to obtain the photoactivable protein probe, as shown in Fig. 1.

**FIG. 1:**
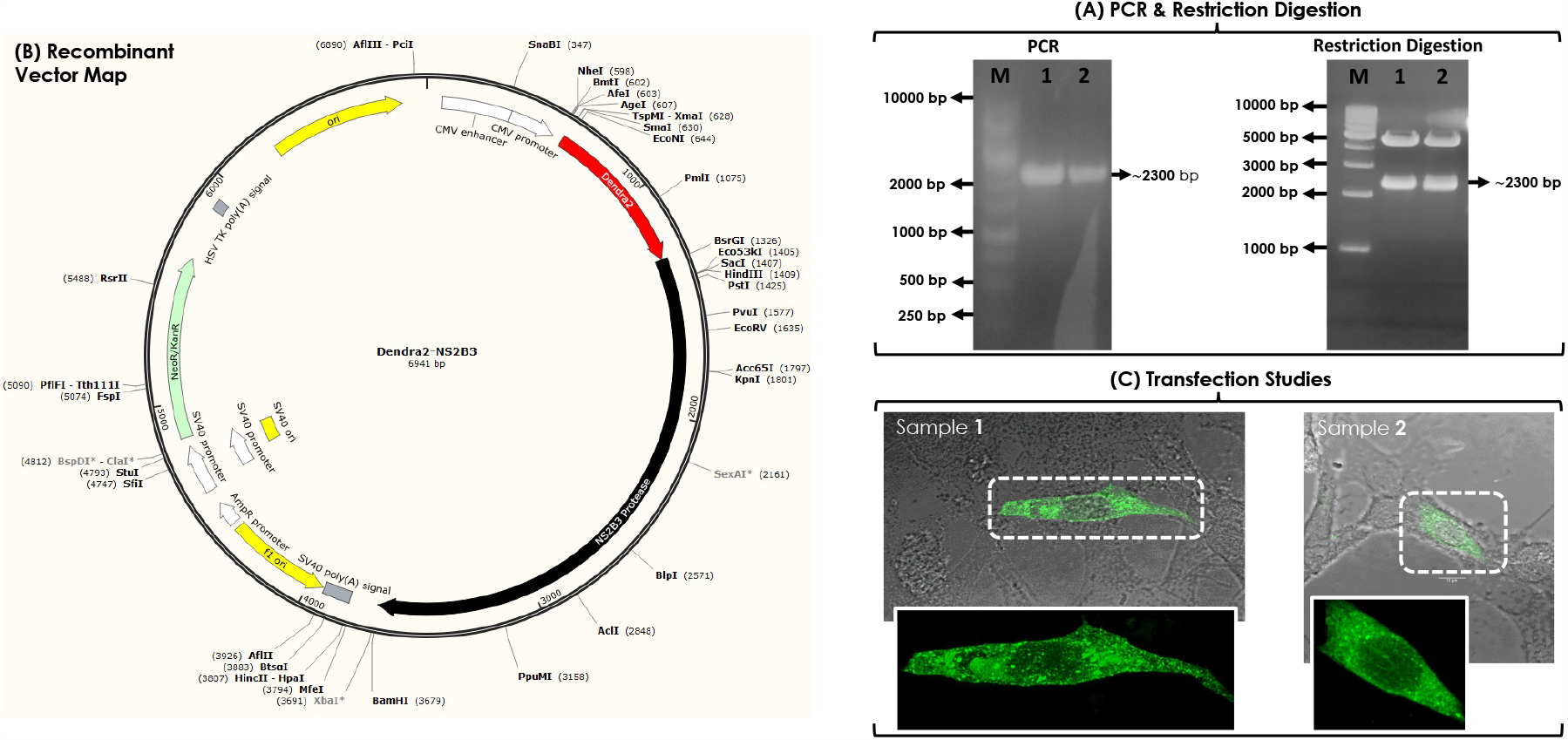
Preparation of photoactivable recombinant plasmid Dendra2-NS2B3: (A) PCR amplification of NS2B3 protease from Dendra2-NS2B3 using specific primers (CMV forward and NS2B3 protease specific reverse). Lane M-1 Kb ladder, Lane 1 and 2 - PCR amplification giving a product around 2300 bp. Restriction analysis Dendra2-NS2B3 releasing NS2B3 protease fragment at Sac I-Bam H1 site. Lane M-1 Kb ladder, Lane 1 and 2-Restriction digestion releasing a product around 2300 bp of NS2B3 protease fragment at Sac I-Bam H1 site. (B) Recombinant plasmid map of Dendra2-NS2B3 with Dendra2 (marked as red) and NS2B3 (marked as black). (C) Validation of recombinant photoactivable plasmid Dendra2-NS2B3 using confocal microscopy where transfected cells shows fluorescence.

The presence of NS2B3 fragment in Dendra2-NS2B3 was confirmed by PCR and restriction digestion analysis. PCR of Dendra2-NS2B3 using NS2B3-specific primers gave an amplicon around 2300bp and restriction analysis using the restriction enzyme pair Sac 1 and Bam H1 released a fragment of 2300 bp along with 1Kb ladder (Fig. 1A). The recombinant nature of Dendra2-NS2B3 was further confirmed by sequencing. Sequence analysis confirmed the intactness of the reading frame while fusing NS2B3 with the fluorophore (Dendra2). The Plasmid map of recombinant plasmid Dendra2-NS2B3 is given in Fig.1B.

The recombinant plasmid Dendra2-NS2B3 was used to transfect NIH3T3 cells (mouse embryonic fibroblast cell line) to study the effect of dengue NS2B3 protease in the cellular system. Fig.1C shows NIH 3T3 cells transfected with recombinant plasmid Dendra 2-NS2B3 followed by incubation for 24h. The transfection was confirmed by exposing it to blue light (470-490 nm) and observing the fluorescence in green (with peak at 507 nm). A strong fluorescence can be seen in the transfected cell along with non-transfected cells.

### B. Super-resolution microscopy revealed the formation of NS2B3 clusters in transfected Cells

We have used Single-molecule localization microscopy (SMLM) to visualize single NS2B3 proteins and its distribution in a cellular environment. A homemade *scanSMLM* system is built to carry out the study, the details of which can be found in Ref. [12]. First, the system is operated in an epifluorescence mode to identify a transfected cell, as shown in Fig. 2A. The corresponding super-resolved image and an enlarged section of a selected region is shown in Fig. 2B. Clustering of NS2B3 proteins is evident along with a small fraction of unclustered ones, post 24 hrs of transfection. Here the clusters are defined as a maximum NS2B3-NS2B3 distance of 60nm for the nearest molecule within the same cluster. The technique allows the determination of physiologically relevant biophysical parameters such as, the area of clusters, number of molecules (NS2B3) per cluster, and cluster density. The corresponding biophysical parameters and their statistical averages are shown in Fig. 2(C,D). Analysis shows an average of 46 *±* 15 clusters per cell with mean of 133 *±* 29 molecules per cluster. The mean area and density of the cluster were found to be 0.0583 *±* 0.0071*μm*^2^ and 3496 *±* 516 *mol/μm*^2^, respectively. These parameters have a direct impact on the rate of infection at a cellular level and thus give a better understanding of the biological mechanism.

**FIG. 2:**
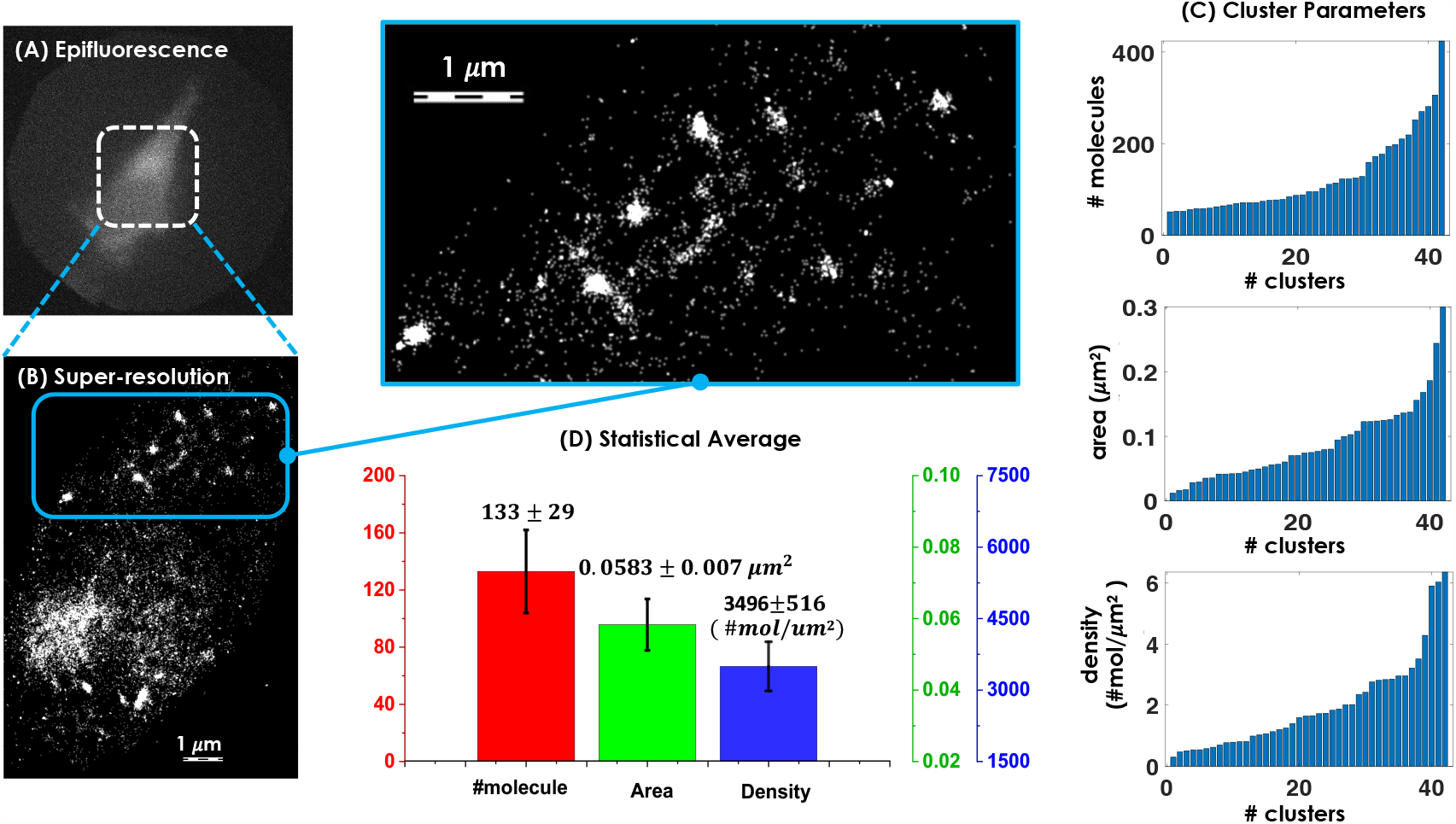
Formation of NS2B3 clusters in transfected NIH3T3 cells: (A) Epifluorescence image of cells transfected with recombinant plasmid Dendra2-NS2B3. (B) Super-resolved image of Dendra2-NS2B3 transfected cells suggests clustering of viral proteins in cellular system. This is further confirmed by enlarged section of selected region. (C) Parameters reveal characteristics of NS2B3 clusters. (D) Statistical analysis estimate an average cluster area of 0.0583 *±* 0.0071*μm*^2^ and cluster density of 3496 *±* 516 *mol/μm*^2^ with an average number of 133 *±* 29 molecules per cluster.

### C. Leflunomide causes dispersion of NS2B3 clusters

Next, we use a drug (leflunomide) to see its effect on NS2B3 clusters. Traditionally, leflunomide is used for the treatment of rheumatoid arthritis which is largely due to its anti-inflammatory and immunomodulatory effects[13] [14]. Moreover, it is known to induce mitochondrial fusion by activating mitofusin 1 and 2 [11]. In addition, we know that NS2B3 acts on mitofusins causing mitochondrial fragmentation [8]. Since both NS2B3 protease and leflunomide are naturally targeted to mitochondria, it encourages the use leflunomide on transfected cells to determine its possible effect on NS2B3 clusters.

To begin with, NIH3T3 cells were transfected with Dendra2-NS2B3 for 24 hrs. Then the cells were washed with PBS for three times and treated with leflunomide for 24 hrs. The drug concentrations used were 250 *nM*, 500 *nM*, 1*μM*, 5*μM*, and 10*μM* . We observed the declustering effect from 250nM onwards compared to control (Dendra2-NS2B3 transfected cells without leflunomide treatment), and it increased at larger concentrations. Declustering indicates the dispersal of NS2B3 clusters from its centre of mass as shown in Fig. 3. Alongside confocal images of sample cells are also shown that indicate strong fluorescence from the transfected cells under study. In addition, we super-resolved an entire 3D cell to understand the collective clustering dynamics using *scanSMLM* as shown in Fig. 4. We have selected 10 layers (planes 1-10) with a *z −* sampling of 500 *nm* in the cell volume beginning from covership, and colormap representation is used to mark the depth. Clear disappearance of NS2B3 clusters is evident in the entire volume with increasing drug concentration. Treatment at low drug concentrations (*≤* 500 *nM*) show breakdown of large clusters into small clusters, whereas a large number of independent NS2B3 molecules are observed at higher concentrations (*≥* 5*μM*), shown a reversal of NS2B3 clustering process. This is indeed the first reported result that promotes the declustering of NS2B3 cluster by the drug leflunomide.

**FIG. 3:**
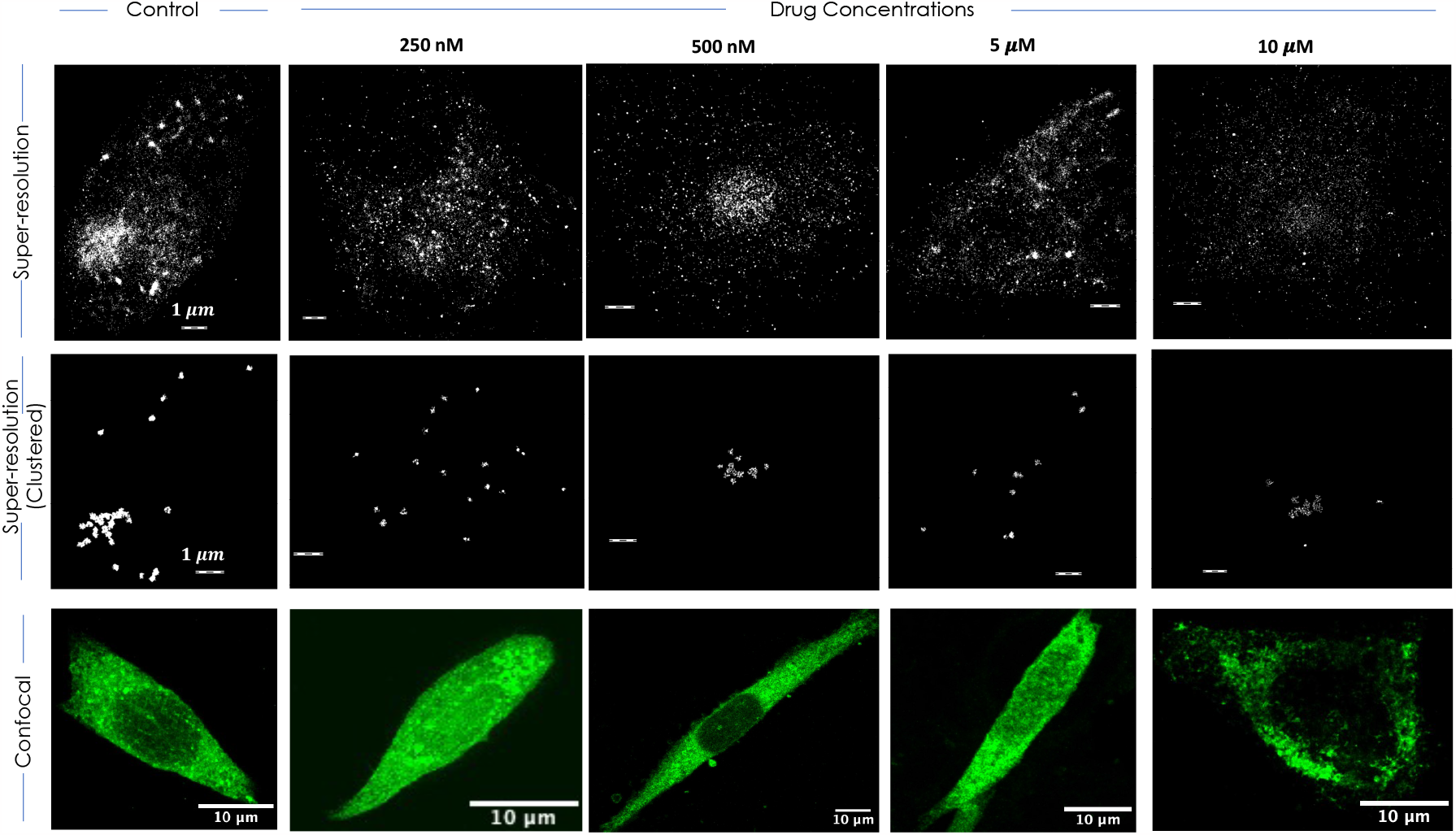
Declustering of NS2B3 clusters upon leflunomide treatment. Single molecule super-resolution study on the viral plasmid Dendr2-NS2B3 transfected cells reveal clustering of viral proteins in the cellular system. Drug treatment study at varying concentration of leflunomide (250 *nM* -10 *μM*) reveal strong declustering of NS2B3 clusters. The third panel shows respective confocal image of transfected cells at different concentrations. Scale bar = 10*μm*

**FIG. 4:**
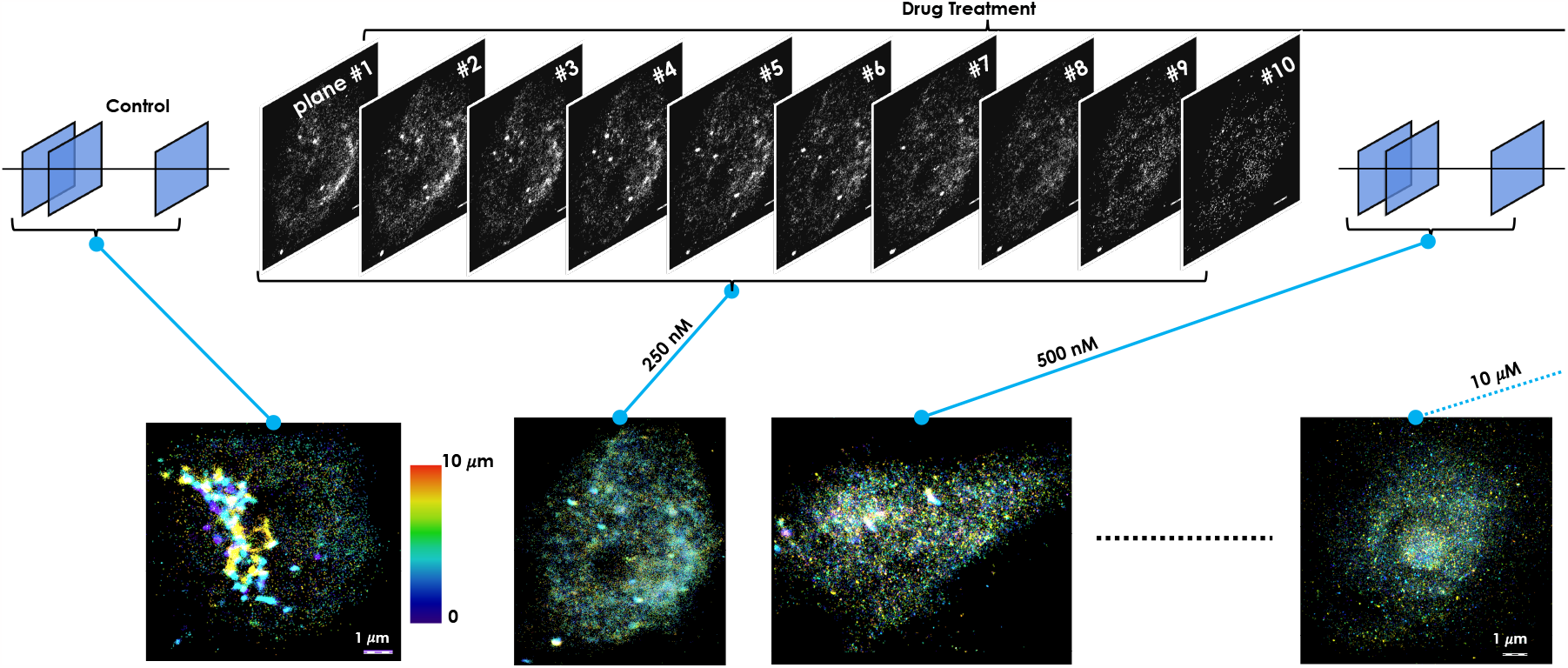
scanSMLM Enabled Volume Imaging of Dendra2-NS2B3 Transfected Cell: Entire 3D imaging a cell showing the action ofleflunomide in transfected NIH3T3 cell. This showing gradual disappearance of clusters in a 3D volume at varying concentration of the drug. For comparison, 3D super-resolved volume of control cell is also shown. The associated colorbar displaying depth information is also shown.

The biophysical parameters related to the formation of clusters, their distribution and size play key roles in the maturation and budding of the viral genome. In this regard, single molecule super-resolution studies enable the determination of critical biophysical parameters such as the number of clusters, percentage of molecules involved in cluster formation, number of molecules/cluster, and mean cluster area (see, Fig. 5). The averages for biophysical parameters are tabulated for control and drug treatment in Table 1. Upon drug treatment, a drastic decrease in the number of clusters is noted overall with increasing concentrations (500 *nM* onwards). However, there is a decrease in the number of clusters in leflunomide-treated cells ranges from 89 *±* 4 to 62 *±* 3 whereas in control it is 133 *±* 29 (Fig. 5). We also observed a decrease in the percentage of NS2B3 molecules engaged in cluster formation when compared to control (53.2 *±* 1.77%) and lowest (10.55 *±* 2.91) at a concentration of 500 nM. In leflunomide-treated cells, the average number of molecules per cluster ranges between 89 *±* 4 to 62 *±* 3, whereas in control it is 133 *±* 29 molecules per cluster. This indicates that large molecules involved in the cluster formation detach due to the declustering action of lefunomide, resulting in smaller clusters. The area occupied by NS2B3 clusters also shows a decreasing trend (see, Fig. 5C). The mean cluster area of the control is found to be 0.0583 *±* 0.0071 *μm*^2^ and it reduced to 0.0266 *±* 0.003*μm*^2^ at a drug concentration of 500 nM. This suggests dispersing nature of leflunomide triggered by either its direct interaction with NS2B3 or indirect interaction involving mitochondria. The exact mechanism is under investigation and may need more extensive studies. Overall, these biophysical parameters collectively point out the fact that leflunomide induces the declustering of NS2B3 clusters in NS2B3 protease expressing cells (Dengue type-2 disease model).

**TABLE I:** **Biophysical parameters related to NS2B3 clustering in cellular system**.

| Parameter | Control | 250 nM | 500 nM | 1 $\mu M$ | 5 $\mu M$ | 10$\mu M$ |
| --- | --- | --- | --- | --- | --- | --- |
| #Clusters | 46±15 | 17±6 | 13±4 | 27±1 | 18±6 | 22±6 |
| #Mol/Cluster | 133±29 | 82±5 | 62±3 | 89±4 | 86±10 | 84±16 |
| Cluster Area ($\mu m^2$) | 0.0583±0.0071 | 0.039±0.004 | 0.0266±0.003 | 0.0299±0.058 | 0.0374±0.005 | 0.0356±0.0076 |
| % of clustered molecules | 53.2±1.77 | 14.89±4.80 | 10.55±2.91 | 23.86±4.56 | 22.09±9.08 | 21.33±9.72 |

**FIG. 5:**
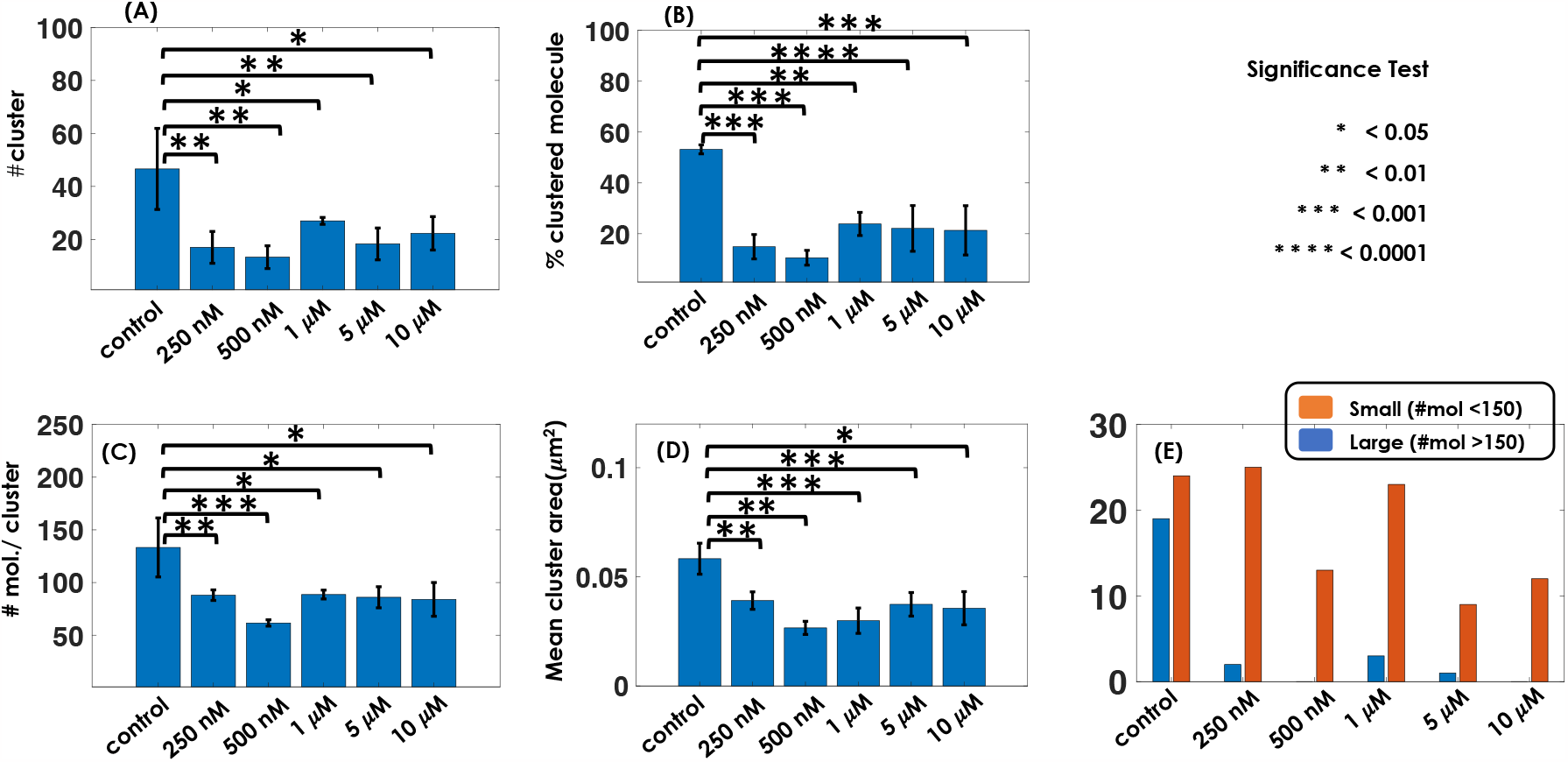
Statistical analysis of cluster parameters indicate declustering of NS2B3 clusters on leflunomide treatment: (A) The number of clusters in leflunomide-treated cells ranges from 27 *±* 1 to 13 *±* 4 whereas in control it is 46 *±* 15.(B) The percentage of molecules engaged in cluster formation is highest in control (53.2 *±* 1.77%) and it decreases with leflunomide treatment (10.55 *±* 2.91% at 500 nM). (C) The average number of molecules/cluster in leflunomide-treated cells ranges between 89 *±* 4 to 62 *±* 3 whereas in control it is 133 *±* 29 molecules per cluster. (D)The mean cluster area of the control is 0.0583 *±* 0.0071*μm*^2^ and the maximum reduction of mean cluster area is at 500 nM of leflunomide (0.0266 *±* 0.003 *μm*^2^). (E) The number of large clusters in control (*∼*18) is high compared to leflunomide-treated ones (*<* 5) whereas more small clusters appeared in leflunomide-treated cells. The statistical significance test carried out between control and treated cells are by *

We summarize the findings related to NS2B3 cluster formation and declustering by leflunomide based on the reported results and ongoing experiments in Fig. 6. In our experiment, lipo-complex containing the plasmid upon transfection enters the cell followed by uncoating of plasmid DNA from lipo complex [15]. It enters the nucleus, and the corresponding mRNA will be synthesized by the process of transcription [16]. Then the mRNA will be carried to the ribosomes located on ER followed by translation and post-translation modifications (expression of NS2B3 protease) [16]. The mature protease is found to form clusters inside the cellular system. Upon treatment with the anti-inflammatory drug leflunomide (through passive diffusion), it enters the cellular system, causing declustering NS2B3 clusters either through direct interaction or via indirect interaction involving other organelles (such as, mitochondria) [17].

**FIG. 6:**
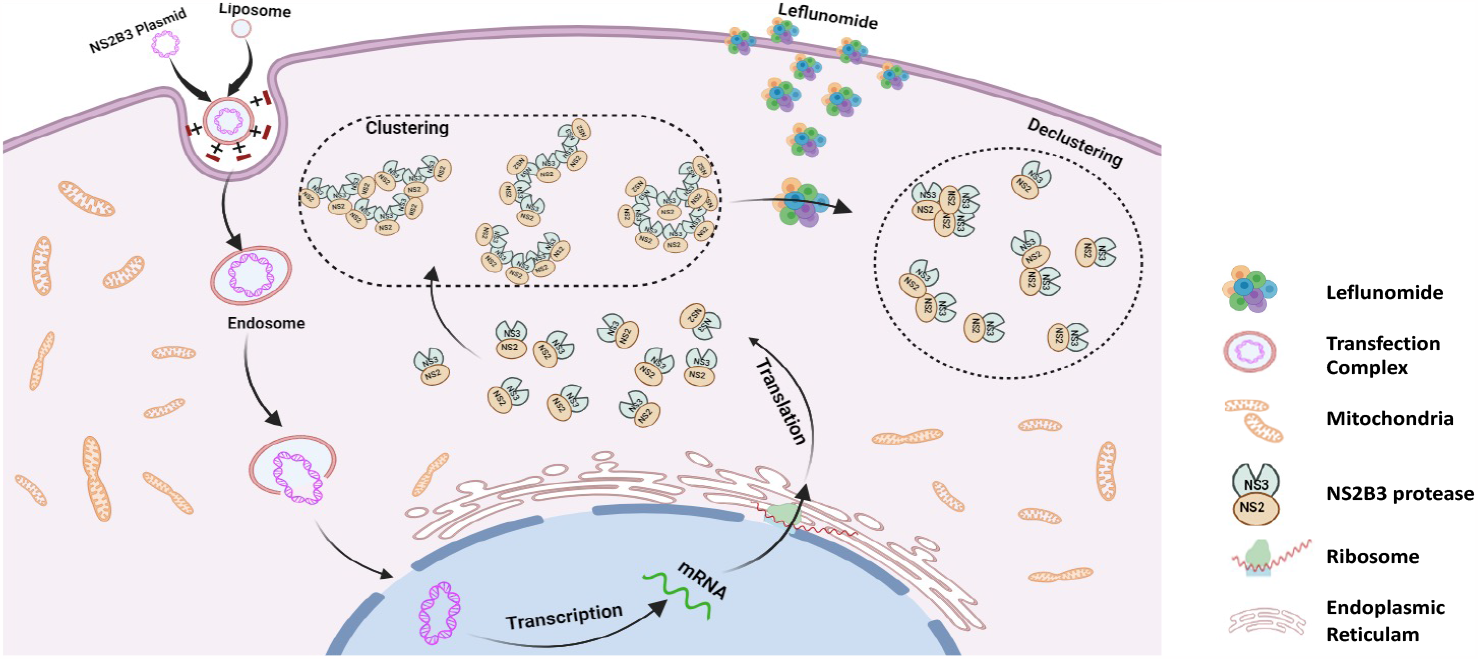
Summary of NS2B3 cluster formation on dengue infection and declustering action of leflunomide in cell: The experimental evidence suggests entry by endocytosis, followed by uncoating, transcription, translation of the viral complex (NS2B3)[16]. The Viral protein complex forms clusters (marked by black dotted circle) in cell. Antiinflammatory drug leflunomide enters the cell to interact with NS2B3 clusters resulting in declustering. All other organelles and proteins are also shown for completeness.

## III. CONCLUSION & DISCUSSION

NS2B3, a dengue viral protease, forms clusters in infected cells. The clustering of viral protein is directly related to its invasion inside the host cell [10]. To investigate, a photoactivable recombinant plasmid (Dendra2-NS2B3) is constructed by standard subclonning, followed by verification using PCR, and restriction digestion. The plasmid is used to transfect NIH3T3 cells, and transfected cells were confirmed using a standard fluorescence microscope. The fixed cell sample was excited using blue light (470 *−* 490 *nm*), and the fluorescence is observed at 507 *nm*. The fluorescence arm is integrated into the *scanSMLM* super-resolution system. This provided a single platform for brightfield, fluorescence, and super-resolution imaging. In the present study, we report the accumulation and clustering of the NS2B3 protease complex in the transfected cells using super-resolution microscopic study (Fig.2). The technique has revealed details of NS2B3 cluster and its characteristics (such as cluster size, #molecules / cluster, fraction of clustered molecules etc) at a single-molecule level. Statistical analysis estimated critical values for the biophysical parameters (average number of clusters of 46 *±* 15, cluster size (Area) 0.0583 *±* 0.0071 *μm*^2^, cluster density of 3496 *±* 516*mol/μm*^2^ and an average of molecules per cluster 133 *±* 29 *mol*.*/cluster*) (Fig.2). It is encouraging to observe the formation of NS2B3 clusters and quantify parameters.

The present study is focused on the effect of the anti-inflammatory drug leflunomide on NS2B3 clusters. The study is performed on a standard model cell line (NIH3T3) that is extensively used for viral study [24] [25]. The cells were transfected with a viral recombinant plasmid Dendra2-NS2B3 (for expressing NS2B3 protein along with fluorophore Dendra2). Post 24 hrs of transfection, the cells were subjected to drug treatment for another 24 hrs, followed by cell fixation. Subsequently, super-resolution studies were carried out using *scanSMLM* system. The study revealed a significant reduction in the size and number of NS2B3 clusters post leflunomide treatment as revealed by standard SMLM (fig.3) and *scanSMLM* (see, Fig. 4). This is well supported statistically (see Fig. 5). Single molecule based super-resolution microscopy has shown the ability to determine physiologically relevant biophysical parameters. Specifically, the study reveals that the number of clusters decreases from 46 *±* 15 to 13 *±* 4, the percentage of molecules engaged in cluster formation falls from 53.2 *±* 1.77% to 10.55 ± 2.91, the average number of molecules per cluster decreases from 133 *±* 29 to 62 *±* 3 and the mean cluster area reduces from 0.0583 *±* 0.0071 *μm*^2^ to 0.0266 *±* 0.003*μm*^2^). These parameters collectively indicate the efficacy of the drug leflunomide in dispersing the clustering of viral protein (NS2B3). In addition, a significant decrease in large clusters is noted at high concentrations (see Fig. 5). Specifically, the cluster disappears at high concentrations as revealed by super-resolution images, and a significant fraction of NS2B3 single molecules is noted (see Fig. 3, and Fig. 4).

Based on the results and experimental evidence in a cellular system, the biophysical processes involved during viral invasion is summarized in Fig. 6. Overall, the reversal of NS2B3 clustering is evident at large drig concentrations. It is obvious that leflunomide intervened with the process of protein clustering. Further investigations are underway to reveal the exact mechanism of declustering by leflunomide and its interaction with NS2B3 viral complex, which may help develop an effective strategy to stop/contain severe dengue infection.

## IV. MATERIALS & METHODS

### A. Clustering Technique and Biophysical Parameter Estimation

Single-molecule clusters are identified using the point-based clustering method. The Euclidean distance between the molecules is found, and if the distance is less than or equal to the cutoff(60nm), then they are assigned to a cluster. This was carried out for all the molecules. After finding all the clusters, a filtering process was carried out to remove unclustered molecules. These unclustered molecules may be biologically insignificant. The number of molecules, Area, Density and percentage of clustered molecules are calculated. This analysis was done for both treated and untreated cells. MATLAB scripts are written for analysis and estimation of biophysical parameters.

### B. Recombinant Plasmids

Recombinant plasmid Dendra2-NS2B3 was generated by subcloning of NS2B3 protease from pcDNA-DENV2-NS2B3, a Gift from Alan Rothman [18] at Sac1-BamH1 site of Dendra2-Tubulin plasmid, a gift from Samuel Hess (University of Maine, USA). NS2B3 protease with Sac1-BamHI overhangs were generated by PCR using NS2B3-specific oligonucleotide primers, NSSacFP with Sac1 site as forward primer and NSBamRP with Bam H1 site as reverse primer. The List and sequence of primers used are given in table II. PCR was performed using Phusion™ High-Fidelity DNA Polymerase (Thermo Fisher Scientific, India). The restriction enzymes used were purchased from Thermo Fisher Scientific, India. Subcloning and transformation were performed using the standard protocol [19]. The recombinant plasmid Dendra2-NS2B3 was used to transfect NIH3T3 cells (mouse embryonic fibroblast cell line) to study the interaction of dengue NS2B3 with the cell organelles.

**TABLE II:**
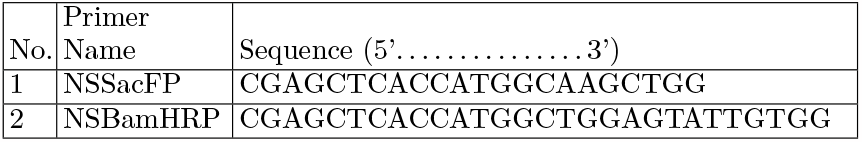
**List of primers used for sub-cloning protein-of-interest (NS2B3) with the photoactivable fluorescent proteins Dendra2**.

### C. Cell Line and Transfection

The recombinant plasmid, Dendra2-NS2B3 used to transfect 3T3 cells (mouse embryonic fibroblast cell line) to study the interaction of dengue NS2B3 with the cellular organelles and its collective dynamics in a cellular system. The cells used in this study were obtained from our collaborator Dr. Upendra Nongthomba (Biological Sciences, Indian Institute of Science, Bangalore, India). Transfection was performed using 1μg of plasmid DNA with Lipofectamine 3000 (Invitrogen, USA) according to manufactures protocol. The cells were cultured on a coverslip (No.0) (Blue star, India) in a 35mm dish with a density of 10^5^ cells/ml. After 12h of incubation, the cells were transfected with Dendra2-NS2B3 plasmids, and the same was used as control. After 24 hrs of post-transfection cells were washed thrice with 1X PBS, and cells were incubated fixed with 3.7% (W/V) paraformaldehyde for 15 min. It was followed by washing with 1X PBS two times. Cover slip was removed from the 35mm dish and placed on a clean glass-slide with mounting media fluorosave (Thermofisher, USA). The Fixed cells were made airproof by applying nail paint on the edges for long time preservation and further investigation.

### D. Drug (leflunomide) treatment

The study is focused on the effect of the anti-inflammatory drug leflunomide (Sigma Aldrich, USA) on NS2B3 induced clusters. The range of drug concentrations used were from 250nM to 10*μM* (250nM, 500nM, 5*μM*, and 10*μM*) on Dendra2-NS2B3 transfected cells. The stock leflunomide (10 mM) was prepared in DMSO. It was diluted to respective working concentrations by diluting in complete medium (DMEM + 10% FBS + 1%Penicillin and Streptomycin).

NIH 3T3 cells were cultured on a coverslip (No.0) (Blue star, India) in a 35mm dish with a density of 1 *×* 10^5^ cells/ml. After 12 h incubation, the cells were transfected with Dendra2-NS2B3 for 24 h. After 24 h of transfection, the cells were treated with respective concentrations of leflunomide for another 24 h. All incubations were at 37^*°*^*C* with 5% CO2. It was followed by fixing and preservation as described in section B. The fixed samples were imaged by confocal and *scanSMLM* super-resolution microscope.

### E. Image acquisition using confocal microscopy and data processing

Fixed samples were imaged in Zeiss LSM 510 inverted microscopy (confocal microscopy Leica SP8, Indian Institute of Science, Bangalore, India) using 63 X oil immersion objective. The cells were illuminated with a 488 nm Laser (argon Multiline), and the emission was captured beyond 500 nm by using a long pass filter and band pass filter and using a 1 Airy unit pin hole. Along with fluorescence image, transmission image is also recorded in a separate detection window.

### F. Super-resolution imaging for Single-molecule analysis

Transfected cells (Dendra2-NS2B3) without leflunomide treatment and transfected cells treated with different concentrations of leflunomide were fixed, and super-resolution studies were carried out using a homemade *scanSMLM* super-resolution system [**?**]. Initially the transfected cells are identified using blue light(470-490 nm), Dendra2-NS2B3 molecules expressed in the transfected cells are activated and excited using laser of wavelength 405 nm and 561 nm respectively. The fluorescence signal was collected on EMCCD(Andor iXon 897) camera with a frame rate of 30Hz. The clusters of single protein molecules (Dendra2-NS2B3) were identified in the reconstructed super-resolved image. Subsequently, biophysical parameters such as, cluster area, number of molecules and density of clusters sere estimated. The effect of leflunomide on cluster formation was quantified using different cluster parameters like the percentage of molecules involved in cluster formation, number of clusters, number of large/small clusters, number of molecules/cluster, and mean cluster area and compared with Dendra2-NS2B3 transfected cells without NS2B3 treatment. The cluster analysis was done using the point-based clustering algorithm [23].

### G. Statistical Analysis

Statistical analysis was carried out using GraphPad Prism software 9. The parameters calculated from cluster analysis are taken for the significant test. Statistical significance test between the control (Dendra2-NS2B3 transfected cells without treatment) and the drug-treated group (Dendra2-NS2B3 transfected cells treated with leflunomide of different concentrations) were carried out using one-way ANOVA. The mean of each parameter corresponding control group was compared with the drug-treated group independently. The obtained p values are less than 0.05 which shows the estimated parameters between control and drug-treated cells are more than 95% statistically different. The corresponding statistical significance (P-values) is indicated as, *∗ P ≤* 0.05, *∗ ∗ P ≤* 0.01, *∗ ∗ ∗ P ≤* 0.001, *∗ ∗ ∗ ∗ P ≤* 0.0001. The statistical analysis is based on a total of 15 cells from 3 separate experiments for each group.

## Acknowledgements

The authors acknowledge ICMR postdoctoral fellowship to JMV. The authors thank Dr. Subhra Mondal (University of Nebraska - Lincoln, USA) for rigorously going through the manuscript and providing valuable inputs.

## Contributions

PPM and JMV conceived the idea. JMV, AS, NP and PPM prepared the samples, carried out the experiments and data analysis. PPM wrote the paper by taking inputs from all the authors.

## Data Availability

The data that support the findings of this study are available from the corresponding author upon request.

